# Arsenic Trioxide Underpins Delayed Neuroinflammation and Impaired Synaptic Integrity involving Integrative Stress Response Signaling

**DOI:** 10.64898/2025.11.30.691457

**Authors:** Jasim Khan, Sazzad Khan, Himanshi Singh, Jianfeng Xiao, Daniel Johnson, Mohammad Athar, Mohammad Moshahid Khan

## Abstract

Arsenic trioxide (ATO), an industrial and environmental chemical with potential for accidental or intentional exposure, exerts profound neurotoxic effects. Its single acute exposure has been linked to delayed neurological and neurodegenerative outcomes. However, the molecular and cellular pathways driving these long-term manifestations remain poorly defined. In this study, combining human iPSC-derived neuronal and *in vivo* mouse model, we uncovered that the acute ATO exposure activates cellular stress pathway, which drives long-term neurotoxic and associated functional outcomes. We found that ATO induces integrated stress response (ISR) signaling in human iPSC-derived neurons. To model the impact of accidental ATO exposure *in vivo*, we administered a single high dose of ATO to C57BL/6J wild type mice. Four weeks post-treatment, ATO-exposed mice displayed neuropsychological symptoms and cognitive deficits. Brain analyses of ATO-challenged mice revealed elevated ISR activity marked by the increased phosphorylation of PERK and eIF2α and upregulation of the transcription factors, CHOP and ATF4. Transcriptomic profiling using bulk RNAseq revealed activation of pathways associated with stress and neuroinflammatory responses. Consistently, increased DNA damage and dysregulation of the STING-mediated innate immune response were also found in the brain of ATO-challenged mice. Pharmacological inhibition of the ISR with a small-molecule inhibitor, ISRIB mitigated ISR activation and preserved synaptic integrity in mouse hippocampal cells. In conclusion, our data identify ISR activation and DNA damage-driven immune dysregulation as key pathogenic drivers of ATO-induced delayed neurotoxicity and cognitive deficits and highlight ISR inhibitors as promising therapeutics to mitigate these effects.

## Introduction

Arsenic trioxide (ATO) is a pervasive environmental and industrial toxicant with profound implications for human health, particularly in situations of accidental exposures or deliberate poisoning. Global production of ATO was estimated at approximately 60,000 metric tons in 2021 (USGS, 2021), with Peru, China, and Morocco emerging as the leading producers in 2023, collectively accounting for nearly 97 % of worldwide output (USGS, 2024). The United States ceased domestic ATO production in 1985 and currently remains entirely reliant on imports to satisfy industrial and pharmaceutical demand. Historical trade data indicate that in 2005, nearly 11,000 metric tons of ATO were imported into the United States, underscoring its dependence on international supply chains. Despite a significant reduction in pesticide-related use following U.S. Environmental Protection Agency regulations since 2021, demand for highly purified ATO continues to grow, reflecting its expanding role in both therapeutic and industrial applications. While ATO serves as a life-saving therapy for acute promyelocytic leukemia (Loh et al. 2024), chronic exposure even at low environmental levels is associated with neurotoxicity, mild cognitive impairment, and an elevated risk of dementia (Gu et al. 2021; Li et al. 2020; O’Bryant et al. 2011; Rashid et al. 2025; Tyler and Allan 2014; Vaidya et al. 2023; Wang et al. 2021). Occupational exposure to high arsenic concentrations has also been linked to behavioral alterations, confusion, and memory loss, underscoring its neurotoxic potential (Morton and Caron 1989; Ratnaike 2003). Experimental studies further link ATO exposure to behavioral deficits, neuroinflammation, oxidative stress, and protein aggregation (Liu et al. 2020; Liu et al. 2024; Nino et al. 2018; Pakzad et al. 2021; Pandey et al. 2025; Zhang et al. 2025), highlighting its paradoxical role as both a therapeutic agent and a potent neurotoxicant.

ATO permeates the blood-brain barrier and accumulates in the central nervous system (CNS), providing a possibility of direct interaction with neural system leading to neuropathogenesis (Itoh et al. 1990; Manthari et al. 2018). Clinical data support this observation through measurable ATO concentrations in cerebrospinal fluid (Kiguchi et al. 2010). Although ATO-induced neurotoxicity is well recognized, the molecular basis of its delayed neurological effects particularly following single large dose exposure remains poorly defined. Emerging evidence implicates oxidative injury, ER stress, and immune dysregulation as converging drivers of neuronal vulnerability across diverse models of environmental toxicant exposure (Chen et al. 2016; Niu et al. 2025; Pan et al. 2024b; Rao et al. 2023; Tam et al. 2020; Watanabe et al. 2017). We and others have shown that arsenic can induce oxidative stress by augmenting mitochondrial damage besides impairing a number of other antioxidant mechanisms (Muzaffar et al. 2023; Nino et al. 2019; Srivastava et al. 2016a; Srivastava et al. 2016b). The integrated stress response (ISR) is an evolutionarily conserved oxidative stress regulated intracellular signaling pathway that controls protein synthesis and protein quality control by reprogramming gene expressions in response to diverse environmental and pathological stressors (Bond et al. 2020; Costa-Mattioli and Walter 2020). Central to this pathway is the eukaryotic translation initiation factor eIF2α, which is robustly phosphorylated at Ser51 under stress conditions. This post translational modification induces a global protein translational block while selectively activating various transcription factors particularly ATF4-dependent transcription, enabling cells to adapt to multiple adverse conditions. ISR activation it is critical in maintaining normal tissue homeostasis under transient stress conditions, however, when prolonged stress remains unresolved, it can trigger inflammatory responses, cell death, and neurodegeneration. Thus, in case of nervous system sustained ISR activation has been implicated in cognitive deficits and compromised neuronal health (Chou et al. 2017; Krukowski et al. 2020b; Morris 2017). Activation of the ISR has been observed following exposure to various environmental toxicants, including ATO, across multiple models of neurotoxicity. This activation frequently coincides with endoplasmic reticulum (ER) stress and induction of the unfolded protein response, key adaptive pathways that converge on eIF2α phosphorylation to restore proteostasis and limit misfolded protein accumulation (Jia et al. 2023; Srivastava et al. 2016b; Wadgaonkar et al. 2022; Weng et al. 2014; Zhang et al. 2015). Protein misfolding is also critical in the pathogenesis of various neurodegenerative disorders (Soto and Estrada 2008; Sweeney et al. 2017). Earlier, a classic inhibitor of ISR, ISRIB was shown to preserve cognitive function, and maintain synaptic integrity, highlighting the involvement of ISR in the models of neurodegeneration (Chou et al. 2017; Hu et al. 2022; Krukowski et al. 2020a; Sidrauski et al. 2013). Although persistent activation of the ISR can inhibit the synthesis of critical DNA repair proteins (Tam et al. 2020), and thereby exacerbates genotoxic stress, the detailed understanding of the mechanism by which ISR orchestrates multiple diverse pathobiology remains unknown. A recent observation demonstrates that ATO induces the accumulation of DNA damage in multiple model systems (Alarifi et al. 2013; Cooper et al. 2022). Additionally, unrepaired damaged DNA can trigger activation of GMP-AMP synthase (cGAS) and stimulator of interferon genes (STING)-mediated innate immune responses, ultimately leading to chronic neuroinflammation and cellular senescence (Ahmad et al. 2024; Dvorkin et al. 2024; Khan et al. 2025a). In chicken and human liver cells, the induction of innate immune responses following arsenic exposure have been reported (Li et al. 2024; Pan et al. 2024a; Pan et al. 2024b). Collectively, these interconnected observations led us to hypothesize that ISR-mediated translational inhibition, impaired DNA repair, and cGAS-STING activation form a deleterious molecular network nexus that may underlie ATO-induced delayed neurotoxicity and progressive cognitive decline.

To investigate the delayed neurotoxic effects of ATO, we established a murine model of single high-dose ATO exposure designed to recapitulate the delayed neurological impairments observed in humans. Our findings reveal that a single high dose of ATO is sufficient to induce delayed long-term consequences and behavioral and cognitive deficits via an interplay of ISR, DNA damage and innate immune signaling.

## Materials and methods

### Cell culture

To assess the impact of ATO on human neurons, iPSC-derived cortical neurons (iXCells Biotechnologies) were cultured on pre-coated plates in cortical neuron maintenance medium (Cat# MD-0093) at 37°C in a COD incubator, following the manufacturer’s instructions. Neurons were exposed to ATO (5 or 10 µM) for 24 hours. Following exposure, cells were fixed and processed for immunocytochemistry to detect ISR activation using phospho-eIF2α (p-eIF2α). Fixed cells were washed twice with 1X phosphate-buffered saline (PBS), permeabilized for 15 minutes with 0.3% Triton X-100 and blocked with 1% bovine serum albumin (BSA) in PBS. Cells were then incubated for 2 hours at room temperature with mouse anti-p-eIF2α (Ser51) or chicken polyclonal MAP2A/B antibody (EnCor Biotechnology Inc; Cat# CPCA-MAP2), followed by incubation with fluorescently conjugated secondary antibodies (Invitrogen, USA) mixed with DAPI for 1 hour. After PBS washes, coverslips were mounted using mounting medium and imaged. HT22 cells were cultured in DMEM supplemented with 10% fetal bovine serum and 1% penicillin-streptomycin at 37°C in a COD incubator. Cells were pretreated with ISRIB (200 nM, 6 h) followed by co-exposure to ATO (5 µM, 24 h) (Li et al. 2025; Sidrauski et al. 2015). After treatment, cells were collected for analysis of ISR signaling and synaptic integrity.

### Mice and treatments

WT (C57BL/6J; Strain #:000664) mice were obtained from Jackson Laboratory, USA. Animals were maintained under standard laboratory conditions with a 12-hour light/dark cycle and provided ad libitum access to commercial rodent chow and water. Following a one-week acclimation period, 3-month-old male WT mice were randomly allocated into several groups and treated once (single dose) orally by gavage with escalated concentrations of ATO (10, and 15 mg/kg) or vehicle. ATO doses were calculated based on reported human fatal ingestion ranges (70-300 mg) (Mochizuki 2019; Ratnaike 2003). The delayed effects of ATO on cognitive and behavioral outcomes were assessed in mice treated with 10 or 15 mg/kg. At the conclusion of behavioral testing, mice were sacrificed, and brain tissues were collected for morphological and biochemical analyses. All animal experiments were conducted in accordance with the guidelines of the Institutional Animal Care and Use Committee and followed the National Institutes of Health’s standards for the care and use of laboratory animals.

### Behavioral Assessments

All behavioral assays, including Morris water maze (MWM), Raised-Beam Task, and Tail suspension tests, were performed as described in our publications (Hori et al. 2022; Ismael et al. 2021; Khan et al. 2018; Khan et al. 2025a; Khan et al. 2025b).

#### Morris Water Maze

Mice were tested in a circular water pool (∼1.5 m diameter, 30 cm depth) maintained at 25-27°C, following previously established methods (Hori et al. 2022; Khan et al. 2018). A hidden platform was initially placed 1 cm above the water surface and made visible with non-toxic white paint for cue training. Mice underwent two consecutive days of cue training, with four trials per day, to familiarize them with the platform’s location. Following cue training, the platform was submerged just below the water surface, and mice were allowed to complete six consecutive days of spatial training, with four trials per day. Escape latency, the time required to locate the platform, was recorded for each trial. Trials were captured using a ceiling-mounted camera, and behavior was analyzed using ANY-maze software (Stoelting Co., USA) to quantify spatial learning, memory retention, and search strategies, providing a robust measure of hippocampal-dependent cognitive function.

#### Raised-Beam Task

Mice were first acclimated to an elevated beam measuring 80 cm in length and 20 mm in width, positioned 50 cm above a padded surface. A 60 W lamp at the starting point served as an aversive stimulus, whereas the opposite end led to a dark escape box. The time taken to traverse the beam and the number of foot slips were recorded for each trial. After initial testing, follow-up tests were performed using additional round and square (9 mm diameter) beams. All testing was conducted in triplicate, and mean values were used for subsequent statistical analyses as previously described (Khan et al. 2018; Khan et al. 2025a).

#### Tail suspension tests

The tail suspension test was conducted to assess depression-like behavior in mice 4 weeks after ATO exposure, as previously described (Khan et al. 2025b). Each mouse was suspended by the tail using adhesive tape from a stable surface, allowing the body to hang freely in a downward-facing position. The test lasted 5 minutes, and behavior was recorded using a video tracking system with automated analysis software (FreezeFrame; Coulbourn, Whitehall, PA, USA). The primary measure was immobility duration, with longer periods of immobility interpreted as indicative of depressive-like behavior. Results are reported as cumulative immobility per minute over the 5-minute testing period.

## RNA sequencing

Total RNA quality and concentration were assessed using the Qubit 4 Fluorometer and Qiagen Qiaxcel system. RNA libraries were prepared with the Illumina Stranded RNA Library Prep Kit and sequenced on an Illumina NextSeq 2000 using a P3 sequencing kit. Raw sequencing reads (FASTQ files) were collected and subjected to quality control using FASTQC. Low-quality bases with a Phred score < Q20 were trimmed before alignment. The processed reads were mapped to the mm38 reference genome using RNA STAR, and the resulting SAM files were used to quantify gene-level read counts. Read counts were normalized across samples using the TMM (Trimmed Mean of M-values) method. Principal component analysis (PCA) and Pearson correlation were performed to assess sample relationships and global transcriptomic profiles. Differential gene expression analysis was conducted using DESeq2, and genes with a p-value ≥ 0.05 or fold change ≤ 1.5 were excluded. The Benjamini-Hochberg false discovery rate (FDR) was applied, retaining genes with FDR < 0.05 as statistically significant. Significant genes were visualized using heatmaps in R, and functional enrichment, including pathway and gene ontology analyses, was performed using STRINGdb.

### Relative quantitative real-time reverse-transcriptase PCR (RT-qPCR) and mitochondrial DNA copy number

Relative mRNA expressions and mitochondrial DNA copy number in mouse tissues were measured using SYBR Green-based RT-qPCR on the Roche LightCycler® 480 System, with primer sequences listed in the supplemental material (**Table S1**). Total RNA was reverse-transcribed into cDNA using random primers and the RETROscript™ Reverse Transcription Kit (ThermoFisher), following the manufacturer’s instructions. GAPDH was used as the internal reference for gene expression analysis. Relative expression and mitochondrial DNA copy number were calculated using the 2^-ΔΔCT^ method and reported as fold change compared with control samples, with genomic DNA used as the reference for mitochondrial DNA quantification.

## Immunofluorescence staining

Immunofluorescent staining was performed as previously described by our group (Khan et al. 2018; Thadathil et al. 2021). Brain sections (25 µm) were prepared using a cryostat and washed with PBS, followed by blocking in 5% BSA with 0.3% Triton X-100. Sections were incubated overnight at 4°C with the following primary antibodies: Iba-1 (1:500; Synaptic Systems) and STING (1:250; ProteinTech). After PBS washes, sections were incubated for 1 hr with Alexa Fluor 555 anti-chicken and Alexa Fluor 488 anti-rabbit secondary antibodies (1:500; Invitrogen) combined with DAPI solution. Sections were then washed and mounted for imaging. Quantification of immunopositive cells was performed by an observer blinded to genotype, using a fluorescent microscope at 40× magnification. Three distinct fields per region were analyzed for each mouse in the hippocampus and cortex, and average cell counts were recorded.

## Western blot analyses

Protein lysates from mouse brain tissue were prepared in ice-cold RIPA lysis buffer (Bio-Rad) supplemented with 1X protease inhibitor cocktail (#I3786, Sigma-Aldrich). Protein concentrations in cell and tissue lysates were determined using a DC Protein Assay kit (Bio-Rad). Protein samples were resolved by SDS-PAGE electrophoresis and subsequently transferred onto a 0.2 µm Immun-Blot PVDF Membrane (Bio-Rad) using Trans-Blot Turbo Transfer System (Bio-Rad). Following protein transfer, membranes were incubated for 1hr at room temperature in blocking buffer (5% nonfat dry milk (Bio-Rad) prepared in a combination of Tris-buffered saline and Tween-20, TBS-T to block nonspecific sites. Membranes were then incubated overnight at 4°C in primary antibodies with dilution suggested by the manufacturer. The following day, membranes were rinsed three times with TBS-T for 10 minutes each on an orbital shaker and then incubated for 2 hr with HRP-conjugated secondary antibodies prepared in 5% nonfat dry milk-TBS-T cocktail. Protein bands were visualized on iBright 1000 Series Image System (ThermoFisher) using enhanced chemiluminescence (Santa Cruz Biotechnology, Dallas, TX, USA). A detailed list of antibodies used in this study is listed in **Table S2.**

## Statistical analysis

All statistical analyses were conducted using GraphPad Prism version 10.2 (GraphPad Software, San Diego, CA). Differences between two groups were evaluated with an unpaired two-tailed t-test, while comparisons across multiple groups were performed using One- or Two-way ANOVA followed by appropriate post hoc tests. Non-parametric behavioral data, such as the number of slips in the raised-beam task, were analyzed using the Mann-Whitney test. Data are presented as mean ± SEM, and a P value of less than 0.05 was considered statistically significant.

## Results

### Arsenic trioxide exposure causes behavioral and neuropsychological symptoms in mice

Exposure to ATO (10 and 15 mg/kg) produces a dose-dependent effects, with higher doses (15 mg/kg) causing greater behavioral deficits and neuropsychological impairments in WT mice. (**Fig. 1 A-D**). ATO-treated WT mice exhibit motor deficits, as evidenced by the increased latency on the 9 mm square and round beam during raised-beam test, including both round and square beams during the raised-beam test (**Fig. 1 A, B**). The Mann-Whitney test reveals that ATO-treated mice exhibit significantly more slips than vehicle-treated mice on both the 9-mm square beam and the 9-mm round beam, indicating compromised coordination and balance. In the MWM test, ATO-treated mice exhibit significantly prolonged latency to locate the hidden platform, indicating impaired spatial learning and memory (**Fig. 1 C**). Furthermore, ATO-treated mice exhibit increased immobility in the tail suspension test (**Fig. 1 D**). In our experiments, ATO-exposed mice display a significantly longer duration of immobility compared with vehicle-treated WT mice (FD,DD = 6.32, P = 0.009) during the 5-minute test period. These findings indicate that ATO exposure induces depressive-like behavior in mice and may contribute to neuropsychological impairments.

**Figure 1.**
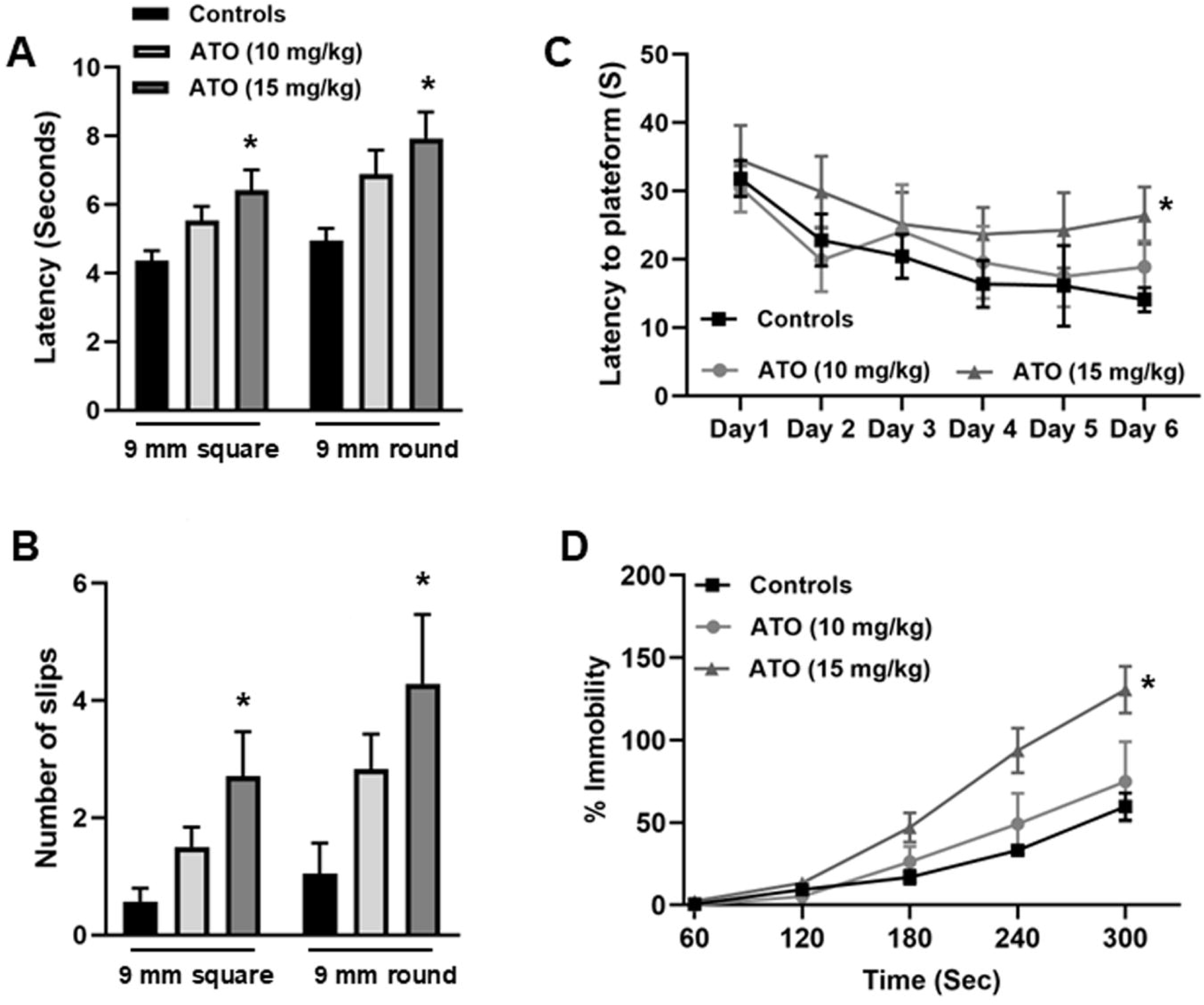
ATO exposure results in cognitive and behavioral deficits. ATO exposure caused significant motor deficits, as evidenced by increased latency to traverse and a higher number of slips. Cognitive impairment was demonstrated by prolonged escape latency in the Morris Water Maze (MWM) test, while depressive-like behavior was indicated by increased immobility time in the tail suspension test (TS). Data were analyzed by one or Two-way ANOVA followed by multiple comparisons tests. Non-parametric behavioral data were evaluated using the Mann-Whitney test. N=6-7 mice/group. The data was presented as mean ± SEM *P < 0.05

Our results are consistent with previous studies reporting ATO-induced affective deficits. (Chang et al. 2015; Liu et al. 2024). Collectively, these results demonstrate that ATO disrupts neuronal circuits involved in motor control, learning, memory, and mood regulation, underscoring its broad neurotoxic effects on the CNS.

### Effect of arsenic trioxide exposure on integrated stress response in the human neurons and the mouse brain

ATO treatment of human iPSC-derived cortical neurons manifests significantly increased phosphorylation of eIF2α at Ser51 in a dose-dependent manner (**Fig. 2 A, B**). This effect mirrors its *in vivo* manifestations, where single acute high-dose ATO exposed WT mice show ISR induction in cortical brain tissue. Specifically, ATO treatment significantly upregulates the accumulation of phospho-PERK and p-eIF2α, along with enhanced expression of ATF4 and CHOP proteins (**Fig. 2 C-G**), the key mediators of stress-induced transcriptional program. These findings demonstrate that acute ATO exposure triggers sustained ISR activation, which may disrupt cellular homeostasis and contribute to long-term neurotoxicity.

**Figure 2.**
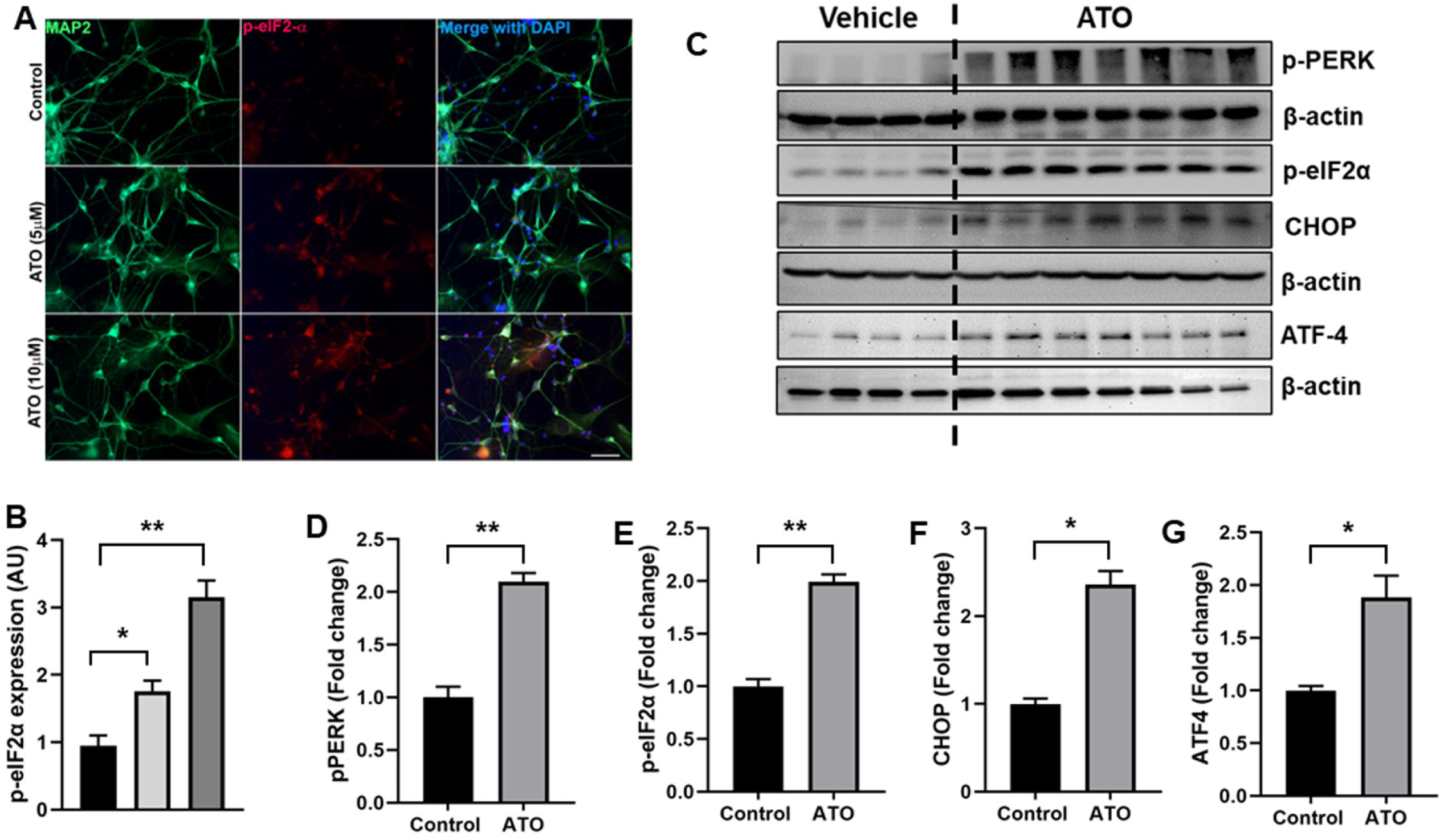
ATO exposure activates the integrated stress response (ISR) in human neurons and mouse brain. (A-B) Representative images of neurons and quantitative analysis showing dose-dependent activation of ISR marker in human iPSC-derived cortical neurons following ATO exposure, as indicated by increased phosphorylation of eIF2α. (C-G) Consistent with in vitro findings, acute ATO exposure in WT mice (N=7) led to robust ISR activation in cortical tissues, as evidenced by increased levels of p-PERK, p-eIF2α, ATF4, and CHOP proteins compared to vehicle controls (N=4). Data were compared between groups using a two-tailed t-test and are expressed as mean ± SEM. *P < 0.05; **P < 0.01.

## Delayed effects of arsenic trioxide on transcriptomics profile in mouse hippocampus

RNA sequencing (RNA-seq) of the hippocampus from WT mice exposed to ATO revealed extensive transcriptional remodeling, with 1,277 differentially expressed genes (DEGs) identified, 409 upregulated and 868 downregulated.

These data indicate a robust and widespread transcriptional response to acute ATO exposure (**Fig. 3A, B**). To better understand the affected biological processes, Gene Ontology (GO) enrichment analysis was performed using STRING-db, identifying 99 significantly enriched pathways (FDR < 0.05). We found that many of these pathways are directly related to cellular stress adaptation and maintenance of tissue homeostasis, including cellular response to stress, protein folding, neuroinflammatory responses, mitochondrial function, and DNA double-strand break repair (**Fig. 3C**). Thus, ATO exposure triggers both proteotoxic and genotoxic stress responses that synergize to manifest neuroinflammation. Heatmap and protein-protein interaction (PPI) analyses reveal that the differentially expressed genes are predominantly involved in stress response pathways (**Fig. S1**) and DNA double-strand break binding (**Fig. S2**), highlighting the activation of proteotoxic and genotoxic stress mechanisms in play in augmenting neurotoxicity of ATO. Molecular analyses of hippocampal tissue corroborate these findings, showing enhanced ISR activation, as evidenced by increased eIF2α and ATF4 levels (**Fig. 3D-F**). In parallel, markers of DNA damage such as γ-H2A.X were elevated (**Fig. 4A, B**), and mitochondrial DNA copy number (**Fig. 4C**) was reduced. Together, these results demonstrate that acute ATO exposure induces persistent transcriptomic alterations that reflect sustained ISR activation, DNA damage, and mitochondrial dysfunction, providing mechanistic insight into the delayed neurotoxic effects of ATO.

**Figure 3.**
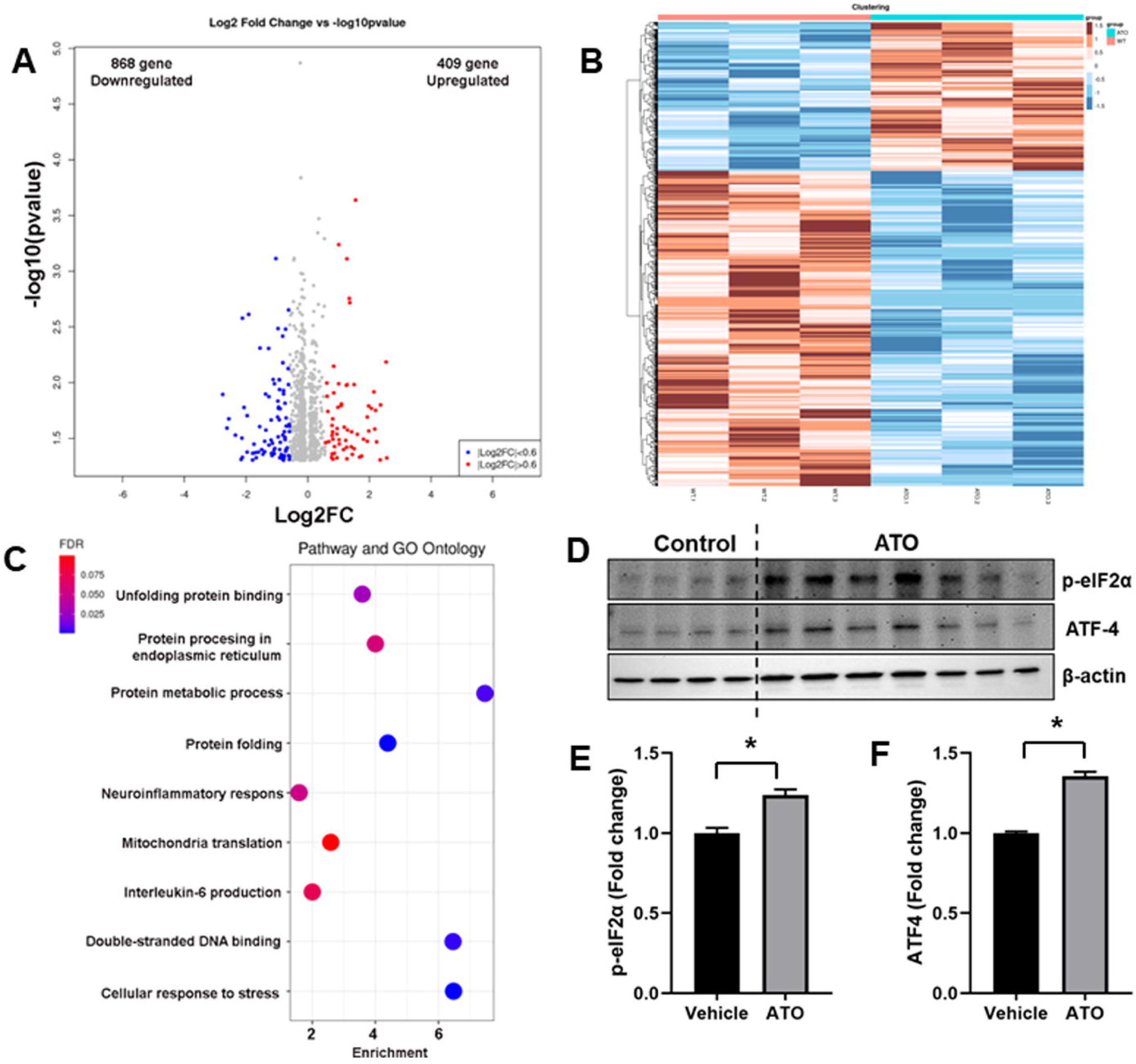
Transcriptomic and molecular signatures of ATO-induced stress responses in WT mouse hippocampus. (A-C) RNA-seq analysis of hippocampal tissue from vehicle and ATO-exposed mice. (A) Volcano plot illustrates significantly upregulated and downregulated genes. (B) Heatmap depicting differentially expressed gene. (C) Bubble plot showing enriched Gene Ontology (GO) pathways, highlighting cellular stress response, unfolded protein binding, and inflammatory signaling. N=3 mice/group. (D-F) Western blot and quantification showing increased phosphorylation of eIF2α and elevated ATF4 expression, confirming activation of ISR in hippocampal tissue. N=4 (vehicle treated) and 7 (ATO treated mice). Data were compared between groups using a two-tailed t-test and are expressed as mean ± SEM. *P < 0.05.

**Figure 4.**
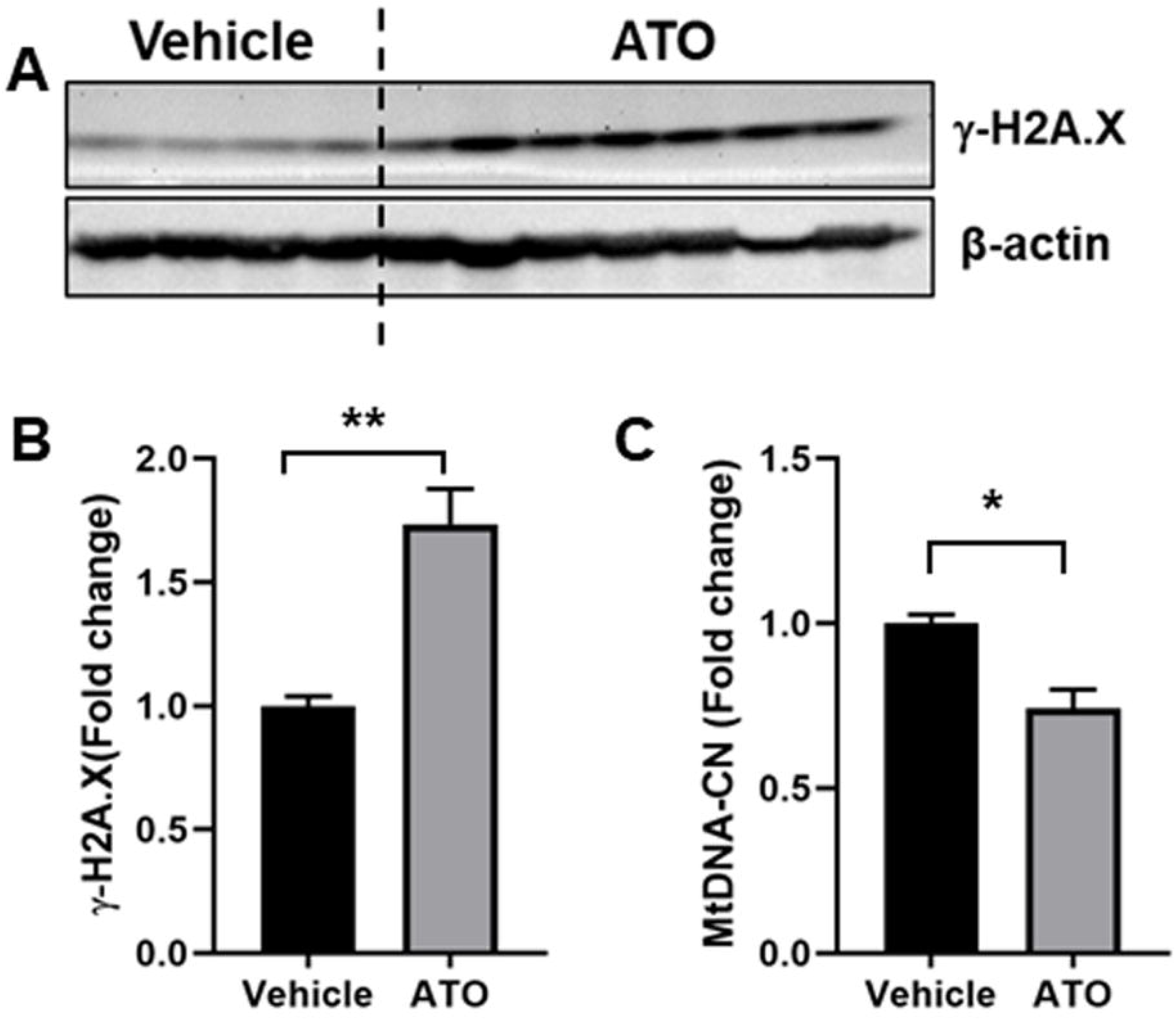
ATO exposure induces DNA damage and mitochondrial dysfunction in mouse brains. (A, B) Representative immunoblots and quantification of γH2A.X levels showing increased DNA damage in the hippocampus of ATO-exposed mice compared with vehicle-treated controls. (C) ATO exposure markedly reduced mitochondrial DNA copy number, indicating mitochondrial impairment. Data were compared between groups using a two-tailed t-test and are expressed as mean ± SEM. *P < 0.05; **P < 0.01.

## Delayed effects of arsenic trioxide on immune responses and neuroinflammation in the brain

Consistent with earlier results and our transcription data analysis, we further examined whether ATO induces STING-dependent immune activation and contributes to chronic neuroinflammation. Our results demonstrate a significant upregulation of STING (**Fig. 5 A, B**). Notably, STING expression was predominantly localized to microglia in the hippocampus, suggesting that microglia are the primary mediators of ATO-induced genotoxic stress (**Fig. 5 F, G**). This activation likely results from the accumulation of cytosolic DNA fragments generated by DNA damage.

**Figure 5.**
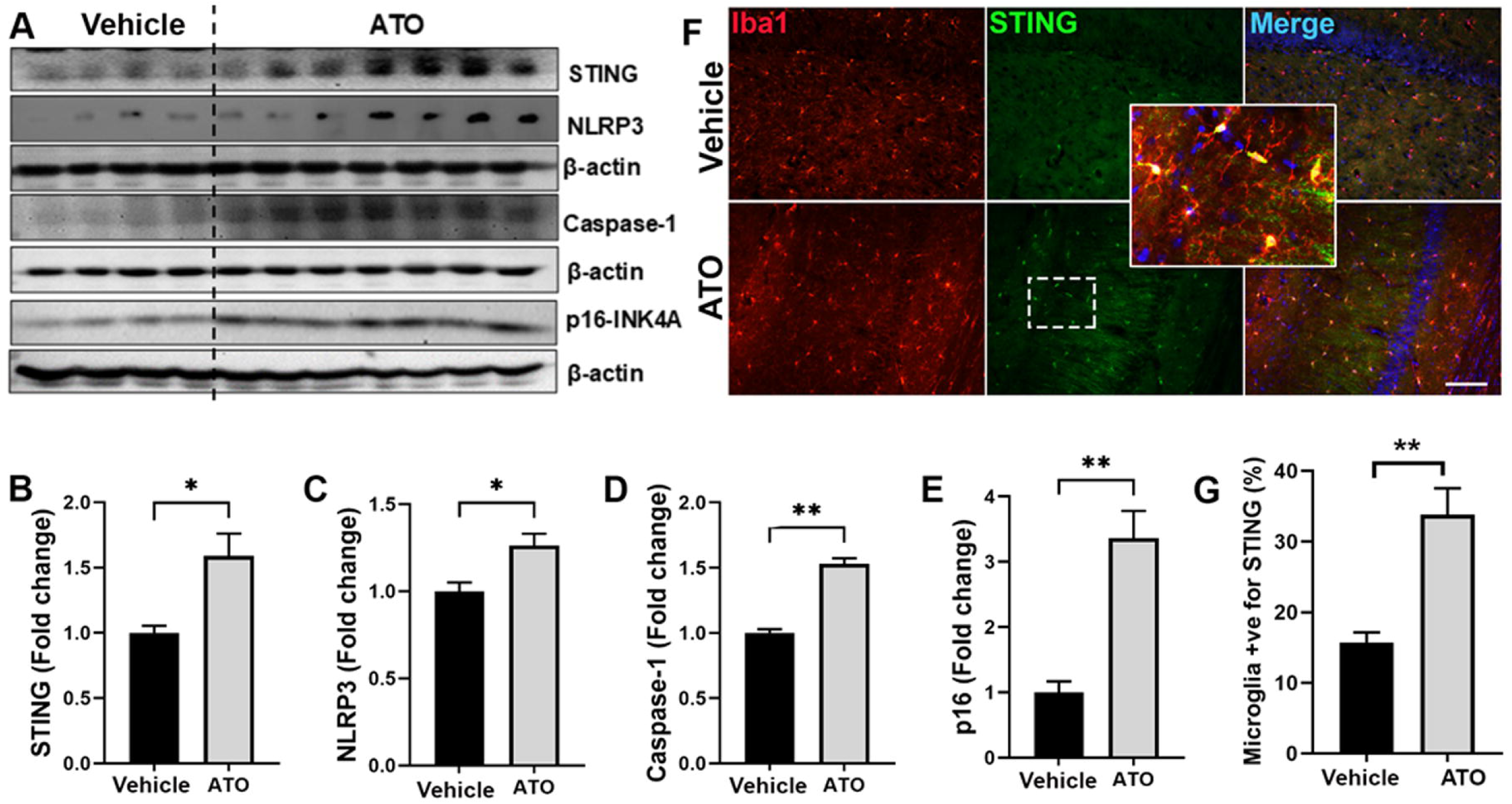
ATO exposure activates STING-mediated neuroinflammation in the hippocampus. (A-E) Representative immunoblots and quantification of STING, NLRP3, caspase-1, and p16 expression in hippocampal tissue of vehicle and ATO-exposed mice, indicating activation of immune signaling and senescence-associated markers. N= 4-7 mice/group. Data were compared between groups using a two-tailed t-test and are expressed as mean ± SEM. *P < 0.05; **P < 0.01. (G-H) Representative immunofluorescence images showing microglia (Iba1, red) and STING (green) in hippocampal sections, with DAPI (blue) labeling nuclei. Quantification of STING-positive microglia demonstrates significant upregulation following ATO exposure. N=3 mice/group. Data are presented as mean ± SEM; **P < 0.01 versus control.

Further analyses revealed increased expression of NLRP3 and caspase-1, indicative of inflammasome activation, a hallmark of robust neuroinflammatory responses **(Fig. 5 C, D)**. Correspondingly, elevated levels of the pro-inflammatory cytokines IL-1β and IL-18, confirm the induction of a pro-inflammatory state, (**Fig. S3**). In addition, we observed upregulation of p16, a marker of cellular senescence **(Fig. 5 E)**, indicating a role of ATO mediated signaling in senescence-associated secretory phenotype. Consistent with our findings in the hippocampus, we also observed upregulation of STING, NLRP3, caspase-1, and p16 in the cortex of WT mice four weeks after single ATO exposure (**Fig. S4**), suggesting a widespread activation of cellular senescence and immune responses across multiple brain regions.

### ISRIB attenuates arsenic trioxide-induced ISR activation and preserves synaptic integrity in hippocampal neurons

Acute exposure to ATO (5 µM, 24 h) triggered robust activation of the ISR in hippocampal HT22 cells, as shown by the elevated levels of ATF4 and CHOP (**Fig. 6A, B**). This sustained stress signaling coincided with a pronounced reduction in postsynaptic density protein 95 (PSD95), a key scaffold critical for synaptic architecture and function (**Fig. 6C**), suggesting an important association of ISR activation and compromised synaptic integrity. Pharmacological inhibition of the ISR with ISRIB (200 nM, 6 h) markedly diminishes ATO-induced stress signaling and prevented PSD95 loss, thus effectively preserving synaptic structure. Collectively, our study highlights ISR modulation as a promising therapeutic strategy for mitigating cognitive deficits and synaptic impairments associated with environmental toxicant exposure.

**Figure 6.**
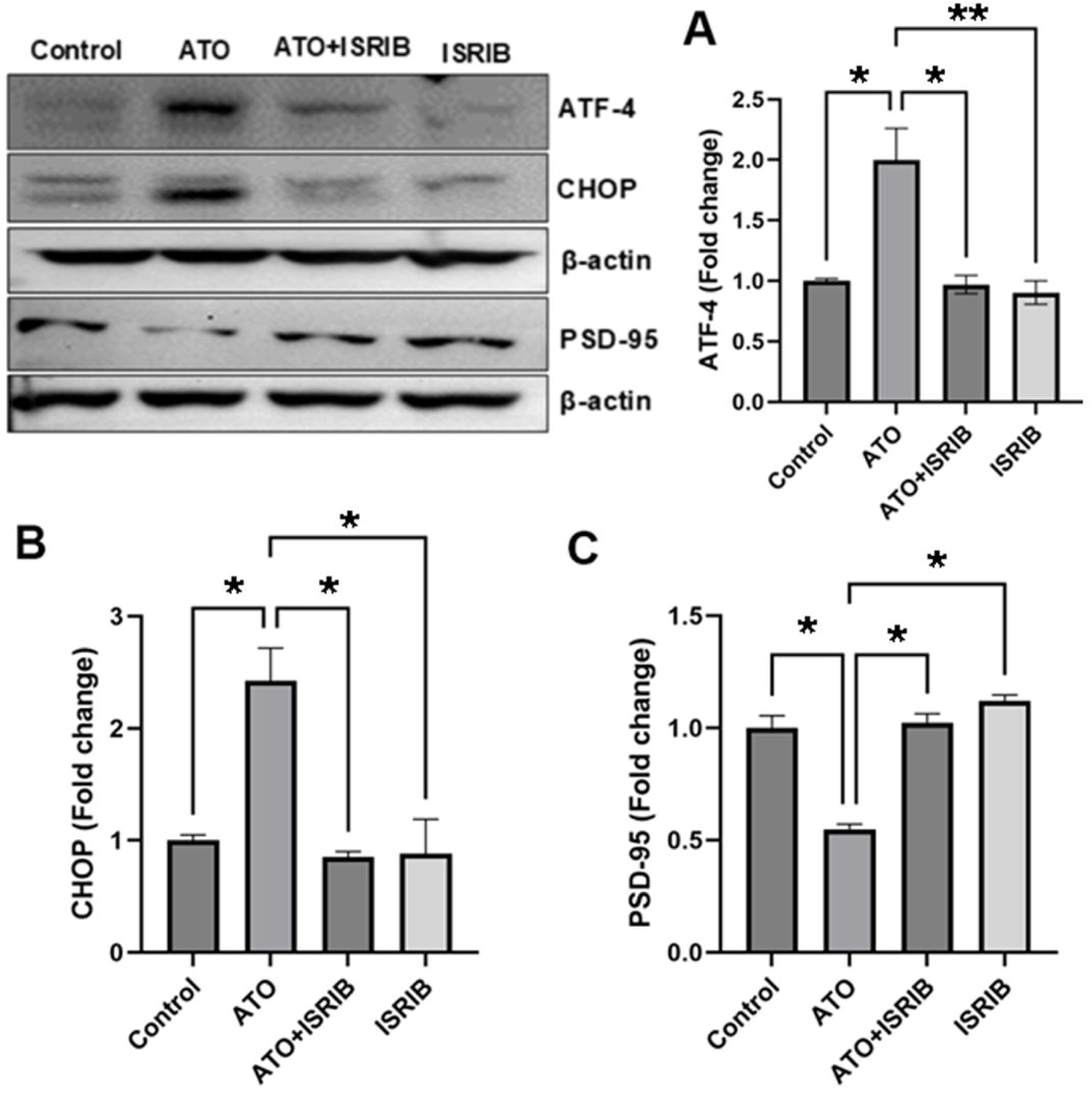
ISRIB counteracts ATO-induced ISR activation and synaptic deficits in hippocampal neurons. (A-C) Representative immunoblots and quantitative analysis showing that ISRIB treatment (200 nM, 6 h) attenuates ATO-induced ISR activation, as evidenced by reduced expression of ATF4 and CHOP, and restores levels of the synaptic scaffold protein PSD95 in hippocampal H22 cells. Data are presented as mean ± SEM; *P < 0.01, **P < 0.001 versus ATO-treated group.s

## Discussion

ATO is a widespread environmental and industrial toxicant, and its exposure poses a significant risk to human health. Here, we show that a single high-dose ATO exposure induces delayed neurotoxic effects, including impairments in cognition, and behavioral functions. Using human neurons, mouse models and unbiased transcriptomics data analysis, we show that ATO drives aberrant ISR signaling, while simultaneously promoting STING-mediated chronic neuroinflammation. These responses collectively disrupt neuronal homeostasis and synaptic integrity. Importantly, pharmacological inhibition of ISR using its classic inhibitor ISRIB mitigates stress signaling and preserves synaptic structure, which not only establishes a mechanistic framework linking acute ATO exposure to delayed neurotoxicity, but also highlights a generalized mechanistic underpinning by which environmental toxicants can drive long-term neurodegenerative-like outcomes.

ATO exposure has long been recognized to induce complex toxico-pathological outcomes across multiple organ systems (Balali-Mood et al. 2025; Muzaffar et al. 2023; Saggu et al. 2024; Tian et al. 2025), while the central nervous system emerges as a critically important vulnerable target (Chu et al. 2025; Gan et al. 2023; Mao et al. 2016; Negi et al. 2024). The pleiotropic nature of ATO toxicity arises from its ability to disrupt a wide range of cellular processes, including redox balance, mitochondrial function, and proteostasis (Chu et al. 2025; Gan et al. 2023; Mao et al. 2016; Negi et al. 2024; Weng et al. 2014) which is also consistent with the results of our study including that our transcriptomics data. One of the critical pathways is the ISR (Iurlaro and Munoz-Pinedo 2016; Kania et al. 2015; Tian et al. 2021), we also previously described. Earlier, chemical exposure has been reported to cause and promote prolonged ISR activation, resulting in maladaptive health outcomes via increased apoptosis or cellular senescence (Bravo-Jimenez et al. 2025; Kalinin et al. 2023) Under stress conditions, protein kinase RNA-like endoplasmic reticulum kinase (PERK) phosphorylates eIF2α, leading to global translational attenuation while selectively promoting the synthesis of stress-related transcription factors such as ATF4 and CHOP (Bell et al. 2016; Wang et al. 2020). Consistently, our findings confirm that ATO exposure robustly activates the ISR in both mouse and human neurons, which is associated with the elevated phosphorylation of PERK and eIF2α and is accompanied by a marked upregulation of the downstream transcriptional regulators ATF4 and CHOP. Clearly, as expected, these molecular alterations underpin neuroinflammation and initiate apoptotic pathways if neural stress is not resolved in timely manner.

Unbiased RNA-seq analysis of the hippocampal tissues revealed that ATO exposure significantly enriches pathways linked to cellular stress responses, unfolded protein binding, and ER-associated protein processing, consistent with our observations of ISR activation. This transcriptional signature is consistent with prior work showing that arsenic trigger ER stress and the unfolded protein response, activating upstream sensors and downstream ISR components (Muzaffar et al. 2023; Srivastava et al. 2016a; Wadgaonkar and Chen 2021). Beyond proteostasis, the RNA-seq profile revealed marked disturbances in mitochondrial-associated pathways. Such mitochondrial vulnerability echoes reports that arsenic perturbs oxidative phosphorylation, disrupts organelle morphology and promotes excessive production of reactive oxygen species (Lu et al. 2014; Srivastava et al. 2016a). These deficits can synergize with ER stress to intensify cellular strain, particularly in neurons, whose high metabolic demands amplify the consequences of impaired energy metabolism. Concurrently, transcripts associated with DNA double-strand binding and genome maintenance were dysregulated, consistent with the documented ability of ATO to induce DNA breaks and interfere with DNA repair machinery, ultimately fostering genomic instability (Kumar et al. 2014; Rao et al. 2023; Tam et al. 2020). In agreement with these transcriptional shifts, we observed increased DNA damage and a reduction in mitochondrial DNA copy number in the brains of ATO-exposed mice. Further, the activation of inflammatory signaling pathways places these molecular disturbances within a broader neuroimmune context. Persistent ER stress, mitochondrial dysfunction and DNA damage are all capable of initiating inflammatory cascades, and accumulating evidence implicates neuroinflammation as a key mediator of cognitive and behavioral impairments caused by environmental toxicants (Andrew and Lein 2021; Guignet et al. 2020). Notably, while ATO has been shown to activate immune responses in non-CNS tissues (Li et al. 2024; Pan et al. 2024b), our findings extend this mechanism to the brain, revealing robust activation pattern of the STING pathway following ATO exposure. Our observation that increased NLRP3, and caspase-1, along with elevated levels of pro-inflammatory cytokines IL-1β and IL-18, which are accompanied by cGAS-STING activation provide a pathogenic signal for sustained neuroinflammation. This is further confirmed by our immunohistochemical analysis identifying microglia within the hippocampus and cortex, the resident immune cells causing ATO-induced neuroinflammation. The cGAS-STING axis serves as a cytosolic DNA-sensing mechanism that bridges genomic instability to innate immune activation and has been increasingly implicated in neurodegenerative disorders (Gulen et al. 2023; Khan et al. 2025a). Our data suggest that ATO-induced DNA damage leads to the accumulation of cytosolic DNA fragments, which engage cGAS, activate STING, and trigger NLRP3 inflammasome signaling. This cascade amplifies neuroinflammation and promotes a feed-forward loop of cellular stress and immune activation. Consistent with prior studies in peripheral tissues (Gan et al. 2023; Li et al. 2024; Pan et al. 2024b) showing ATO-induced cGAS-STING and NLRP3 activation, our results highlight that similar mechanisms operate in the brain, linking genotoxic stress to chronic neuroinflammation and neuronal dysfunction. We also observed increased p16 expression in the brains of ATO-exposed mice, suggesting that ATO-induced stress promotes a senescence-like state in the CNS. p16 is a well-established marker of cellular senescence, functioning as a cyclin-dependent kinase inhibitor that enforces irreversible cell cycle arrest in response to DNA damage or chronic stress (Gonzalez-Gualda et al. 2021; Khan et al. 2025a; Shreeya et al. 2023). Although exact cell-type specificity was not assessed, its upregulation, together with activation of the cGAS-STING pathway, points to a maladaptive shift toward chronic neuroinflammation and irreversible cellular dysfunction, a hallmark of delayed neurotoxicity.

Our behavioral studies are in coordination with our molecular data and further indicate the delayed neurotoxic effects of ATO. In this regard, motor deficits in the raised-beam tasks, as well as cognitive impairments in the MWM test are important indicators of these delayed toxic manifestations. ATO-treated mice exhibit prolonged latency to reach the hidden platform and spend less time in the target quadrant, indicating impaired spatial learning and memory. Previous studies demonstrating that arsenic exposure leads to deficits in motor coordination and cognitive function in rodents (Mehta et al. 2020; Pandey et al. 2025; Yan et al. 2022) and humans (Gong et al. 2011; O’Bryant et al. 2011; Tyler and Allan 2014; Vaidya et al. 2023) are also consistent with our data.

Additionally, the increased immobility observed during the tail suspension test following ATO treatment implies that ATO may drive behavioral traits consistent with depressive pathology. This is highly likely that these effects are due to targeting ATO on pathways that impact both the immune system and the CNS. Neuroinflammation as is induced by ATO, has been shown to be a central player in the pathophysiology of mood disorders. Pro-inflammatory cytokines are known interrupters of signals in neurotransmission and thus contribute to depressive-like behaviors (Liu et al. 2024).

ATO accumulates in the brain particularly in the cortex and hippocampus, which is consistent with its ability to compromise and cross the blood brain barrier (Itoh et al., 1990, Watanabe et al., 2009, Manthari et al., 2018). Here we show that in hippocampal HT22 neurons, ATO triggers robust ISR activation, as reflected by elevated ATF4 and CHOP expression, coupled with a marked reduction in PSD95, a postsynaptic scaffolding protein critical for maintaining synaptic architecture and plasticity. This pattern underscores a direct link between stress pathway overactivation and synaptic degradation, consistent with previous reports implicating maladaptive ISR signaling in neurodegenerative processes (Bravo-Jimenez et al. 2025; Costa-Mattioli and Walter 2020). This conclusion is supported by our observations that ISRIB treatment markedly attenuates ATO-induced downregulation of PSD95, and thereby preserving synaptic integrity by reducing ISR. It is known that ISRIB acts by stabilizing the eIF2B complex, restoring translation even in the presence of phosphorylated eIF2α, thus allowing neurons to maintain protein synthesis required for synaptic maintenance and function (Tsai et al. 2018; Zyryanova et al. 2021). These results highlight the functional relevance of ISR hyperactivation as a driver of ATO-induced synaptic pathology and establish ISRIB as a potential neuroprotective agent capable of restoring proteostatic balance and synaptic health. Earlier studies also showed potential of ISRIB to abrogate neurodegenerative effects in various animal models (Chou et al. 2017; Hu et al. 2022; Krukowski et al. 2020a; Sidrauski et al. 2013).

In conclusion, our study uncovers a mechanistic framework for ATO-induced delayed neurotoxicity by augmenting maladaptive ISR activation, DNA damage and STING-driven neuroinflammation. These clastogenic effects disrupt neuronal homeostasis, leading to synaptic loss and behavioral deficits. By linking cellular stress pathways to long-term neurobehavioral outcomes, these findings provide critical insight into the plausible therapeutic benefit of ISRIB for restoring or delaying appearance of neural toxicity. Although ISRIB in this *in vitro* study is tested to effectively attenuate ISR signaling and preserved synaptic integrity, future studies are needed to confirm its neuroprotective efficacy *in vivo* and to define the temporal and spatial dynamics of ATO accumulation and disposition in the brain. Collectively, this work advances our understanding of delayed neurotoxicity of acute single high dose ATO exposure as may occur during warfare or terrorist or accidental industrial blast. Our data also identifies a potential therapeutic strategy to counteract its enduring neurological impact. Furthermore, ISRIB, despite its unfavorable pharmacological profile, remains a potentially powerful and non-toxic medical countermeasure to block the neural toxicity of ATO.

## Data availability

The data generated and analyzed in this study are available from the corresponding author upon reasonable request.

## Statements and Declarations

The authors declare no conflicts of interest.

## Acknowledgement.

This work was partly supported by the NIH grants U01NS118095, R21 NS128519, Alzheimer’s Association AARG-NTF-22-972518 and the Department of Neurology internal fund.

## References

1. Ahmad A, Braden A, Khan S, Xiao J, Khan MM (2024) Crosstalk between the DNA damage response and cellular senescence drives aging and age-related diseases. Semin Immunopathol 46(3-4):10 doi:10.1007/s00281-024-01016-7

2. Alarifi S, Ali D, Alkahtani S, Siddiqui MA, Ali BA (2013) Arsenic trioxide-mediated oxidative stress and genotoxicity in human hepatocellular carcinoma cells. Onco Targets Ther 6:75–84 doi:10.2147/OTT.S38227

3. Andrew PM, Lein PJ (2021) Neuroinflammation as a Therapeutic Target for Mitigating the Long-Term Consequences of Acute Organophosphate Intoxication. Front Pharmacol 12:674325 doi:10.3389/fphar.2021.674325

4. Balali-Mood M, Eizadi-Mood N, Hassanian-Moghaddam H, et al. (2025) Recent advances in the clinical management of intoxication by five heavy metals: Mercury, lead, chromium, cadmium and arsenic. Heliyon 11(4):e42696 doi:10.1016/j.heliyon.2025.e42696

5. Bell MC, Meier SE, Ingram AL, Abisambra JF (2016) PERK-opathies: An Endoplasmic Reticulum Stress Mechanism Underlying Neurodegeneration. Curr Alzheimer Res 13(2):150–63 doi:10.2174/1567205013666151218145431

6. Bond S, Lopez-Lloreda C, Gannon PJ, Akay-Espinoza C, Jordan-Sciutto KL (2020) The Integrated Stress Response and Phosphorylated Eukaryotic Initiation Factor 2alpha in Neurodegeneration. J Neuropathol Exp Neurol 79(2):123–143 doi:10.1093/jnen/nlz129

7. Bravo-Jimenez MA, Sharma S, Karimi-Abdolrezaee S (2025) The integrated stress response in neurodegenerative diseases. Mol Neurodegener 20(1):20 doi:10.1186/s13024-025-00811-6

8. Chang CY, Guo HR, Tsai WC, et al. (2015) Subchronic Arsenic Exposure Induces Anxiety-Like Behaviors in Normal Mice and Enhances Depression-Like Behaviors in the Chemically Induced Mouse Model of Depression. Biomed Res Int 2015:159015 doi:10.1155/2015/159015

9. Chen G, Mao J, Zhao J, et al. (2016) Arsenic trioxide mediates HAPI microglia inflammatory response and the secretion of inflammatory cytokine IL-6 via Akt/NF-kappaB signaling pathway. Regul Toxicol Pharmacol 81:480–488 doi:10.1016/j.yrtph.2016.09.027

10. Chou A, Krukowski K, Jopson T, et al. (2017) Inhibition of the integrated stress response reverses cognitive deficits after traumatic brain injury. Proc Natl Acad Sci U S A 114(31):E6420–E6426 doi:10.1073/pnas.1707661114

11. Chu X, Li C, Hao Y, et al. (2025) Targeting Nrf2/HO-1 signaling by crocin: Role in attenuation of arsenic trioxide-induced neurotoxicity in mice. J Ethnopharmacol 337(Pt 2):118858 doi:10.1016/j.jep.2024.118858

12. Cooper KL, Liu R, Zhou X (2022) Particulate arsenic trioxide induces higher DNA damage and reactive oxygen species than soluble arsenite in lung epithelial cells. Toxicol Appl Pharmacol 457:116320 doi:10.1016/j.taap.2022.116320

13. Costa-Mattioli M, Walter P (2020) The integrated stress response: From mechanism to disease. Science 368(6489) doi:10.1126/science.aat5314

14. Dvorkin S, Cambier S, Volkman HE, Stetson DB (2024) New frontiers in the cGAS-STING intracellular DNA-sensing pathway. Immunity 57(4):718–730 doi:10.1016/j.immuni.2024.02.019

15. Gan R, Liu H, Wu S, et al. (2023) Curcumin Alleviates Arsenic Trioxide-Induced Inflammation and Pyroptosis via the NF-kappaB/NLRP3 Signaling Pathway in the Hypothalamus of Ducks. Biol Trace Elem Res 201(5):2503–2511 doi:10.1007/s12011-022-03321-4

16. Gong G, Hargrave KA, Hobson V, et al. (2011) Low-level groundwater arsenic exposure impacts cognition: a project FRONTIER study. J Environ Health 74(2):16–22

17. Gonzalez-Gualda E, Baker AG, Fruk L, Munoz-Espin D (2021) A guide to assessing cellular senescence in vitro and in vivo. FEBS J 288(1):56–80 doi:10.1111/febs.15570

18. Gu L, Yu J, Fan Y, et al. (2021) The Association Between Trace Elements Exposure and the Cognition in the Elderly in China. Biol Trace Elem Res 199(2):403–412 doi:10.1007/s12011-020-02154-3

19. Guignet M, Dhakal K, Flannery BM, et al. (2020) Persistent behavior deficits, neuroinflammation, and oxidative stress in a rat model of acute organophosphate intoxication. Neurobiol Dis 133:104431 doi:10.1016/j.nbd.2019.03.019

20. Gulen MF, Samson N, Keller A, et al. (2023) cGAS-STING drives ageing-related inflammation and neurodegeneration. Nature 620(7973):374–380 doi:10.1038/s41586-023-06373-1

21. Hori RT, Moshahid Khan M, Xiao J, Hargrove PW, Moss T, LeDoux MS (2022) Behavioral and molecular effects of Ubtf knockout and knockdown in mice. Brain Res 1793:148053 doi:10.1016/j.brainres.2022.148053

22. Hu Z, Yu P, Zhang Y, et al. (2022) Inhibition of the ISR abrogates mGluR5-dependent long-term depression and spatial memory deficits in a rat model of Alzheimer’s disease. Transl Psychiatry 12(1):96 doi:10.1038/s41398-022-01862-9

23. Ismael S, Nasoohi S, Li L, et al. (2021) Thioredoxin interacting protein regulates age-associated neuroinflammation. Neurobiol Dis 156:105399 doi:10.1016/j.nbd.2021.105399

24. Itoh T, Zhang YF, Murai S, et al. (1990) The effect of arsenic trioxide on brain monoamine metabolism and locomotor activity of mice. Toxicol Lett 54(2-3):345–53 doi:10.1016/0378-4274(90)90202-w

25. Iurlaro R, Munoz-Pinedo C (2016) Cell death induced by endoplasmic reticulum stress. FEBS J 283(14):2640–52 doi:10.1111/febs.13598

26. Jia Y, Ma X, He B, et al. (2023) Manganese triggers persistent activation of the integrated stress response by inhibition of SIRT1 on deacetylation of GADD34. Sci Total Environ 887:164124 doi:10.1016/j.scitotenv.2023.164124

27. Kalinin A, Zubkova E, Menshikov M (2023) Integrated Stress Response (ISR) Pathway: Unraveling Its Role in Cellular Senescence. Int J Mol Sci 24(24) doi:10.3390/ijms242417423

28. Kania E, Pajak B, Orzechowski A (2015) Calcium homeostasis and ER stress in control of autophagy in cancer cells. Biomed Res Int 2015:352794 doi:10.1155/2015/352794

29. Khan MM, Xiao J, Patel D, LeDoux MS (2018) DNA damage and neurodegenerative phenotypes in aged Ciz1 null mice. Neurobiol Aging 62:180–190 doi:10.1016/j.neurobiolaging.2017.10.014

30. Khan S, Delotterie DF, Xiao J, et al. (2025a) Crosstalk between DNA damage and cGAS-STING immune pathway drives neuroinflammation and dopaminergic neurodegeneration in Parkinson’s disease. Brain Behav Immun:106065 doi:10.1016/j.bbi.2025.106065

31. Khan S, Singh H, Xiao J, Khan MM (2025b) DNA Damage Response Regulation Alleviates Neuroinflammation in a Mouse Model of alpha-Synucleinopathy. Biomolecules 15(7) doi:10.3390/biom15070907

32. Kiguchi T, Yoshino Y, Yuan B, et al. (2010) Speciation of arsenic trioxide penetrates into cerebrospinal fluid in patients with acute promyelocytic leukemia. Leuk Res 34(3):403–5 doi:10.1016/j.leukres.2009.08.001

33. Krukowski K, Nolan A, Frias ES, et al. (2020a) Small molecule cognitive enhancer reverses age-related memory decline in mice. Elife 9 doi:10.7554/eLife.62048

34. Krukowski K, Nolan A, Frias ES, et al. (2020b) Integrated Stress Response Inhibitor Reverses Sex-Dependent Behavioral and Cell-Specific Deficits after Mild Repetitive Head Trauma. J Neurotrauma 37(11):1370–1380 doi:10.1089/neu.2019.6827

35. Kumar S, Yedjou CG, Tchounwou PB (2014) Arsenic trioxide induces oxidative stress, DNA damage, and mitochondrial pathway of apoptosis in human leukemia (HL-60) cells. J Exp Clin Cancer Res 33(1):42 doi:10.1186/1756-9966-33-42

36. Li Q, Zhang C, Qi E, et al. (2025) ISRIB facilitates post-spinal cord injury recovery through attenuation of neuronal apoptosis and modulation of neuroinflammation. J Orthop Translat 51:119–131 doi:10.1016/j.jot.2025.01.003

37. Li X, Pan YF, Chen YB, et al. (2024) Arsenic trioxide augments immunogenic cell death and induces cGAS-STING-IFN pathway activation in hepatocellular carcinoma. Cell Death Dis 15(4):300 doi:10.1038/s41419-024-06685-8

38. Li XL, Zhan RQ, Zheng W, Jiang H, Zhang DF, Shen XL (2020) Positive association between soil arsenic concentration and mortality from alzheimer’s disease in mainland China. J Trace Elem Med Biol 59:126452 doi:10.1016/j.jtemb.2020.126452

39. Liu P, Xue Y, Zheng B, et al. (2020) Crocetin attenuates the oxidative stress, inflammation and apoptosisin arsenic trioxide-induced nephrotoxic rats: Implication of PI3K/AKT pathway. Int Immunopharmacol 88:106959 doi:10.1016/j.intimp.2020.106959

40. Liu S, Lei T, Wang L, et al. (2024) Taurine Reverses Arsenic-Induced Inhibition of Hippocampal Neurogenesis and Depression-Like Behavior in Mice. J Biochem Mol Toxicol 38(11):e70037 doi:10.1002/jbt.70037

41. Loh Z, Ashby M, Van Veldhuizen E, et al. (2024) Arsenic-induced neurotoxicity in patients with acute promyelocytic leukaemia. Br J Haematol 204(5):1732–1739 doi:10.1111/bjh.19297

42. Lu TH, Tseng TJ, Su CC, et al. (2014) Arsenic induces reactive oxygen species-caused neuronal cell apoptosis through JNK/ERK-mediated mitochondria-dependent and GRP 78/CHOP-regulated pathways. Toxicol Lett 224(1):130–40 doi:10.1016/j.toxlet.2013.10.013

43. Manthari RK, Tikka C, Ommati MM, et al. (2018) Arsenic induces autophagy in developmental mouse cerebral cortex and hippocampus by inhibiting PI3K/Akt/mTOR signaling pathway: involvement of blood-brain barrier’s tight junction proteins. Arch Toxicol 92(11):3255–3275 doi:10.1007/s00204-018-2304-y

44. Mao J, Yang J, Zhang Y, et al. (2016) Arsenic trioxide mediates HAPI microglia inflammatory response and subsequent neuron apoptosis through p38/JNK MAPK/STAT3 pathway. Toxicol Appl Pharmacol 303:79–89 doi:10.1016/j.taap.2016.05.003

45. Mehta K, Kaur B, Pandey KK, Kaler S, Dhar P (2020) Curcumin supplementation shows modulatory influence on functional and morphological features of hippocampus in mice subjected to arsenic trioxide exposure. Anat Cell Biol 53(3):355–365 doi:10.5115/acb.18.169

46. Mochizuki H (2019) Arsenic Neurotoxicity in Humans. Int J Mol Sci 20(14) doi:10.3390/ijms20143418

47. Morris A (2017) Neuroendocrinology: Integrated stress response linked to TBI. Nat Rev Endocrinol 13(9):501 doi:10.1038/nrendo.2017.102

48. Morton WE, Caron GA (1989) Encephalopathy: an uncommon manifestation of workplace arsenic poisoning? Am J Ind Med 15(1):1–5 doi:10.1002/ajim.4700150102

49. Muzaffar S, Khan J, Srivastava R, Gorbatyuk MS, Athar M (2023) Mechanistic understanding of the toxic effects of arsenic and warfare arsenicals on human health and environment. Cell Biol Toxicol 39(1):85–110 doi:10.1007/s10565-022-09710-8

50. Negi V, Singh P, Singh L, Pandey RK, Kumar S (2024) A Comprehensive Review on Molecular Mechanism Involved in Arsenic Trioxide Mediated Cerebral Neurodegenerative and Infectious Diseases. Infect Disord Drug Targets 24(3):e131123223549 doi:10.2174/0118715265262440231103094609

51. Nino SA, Martel-Gallegos G, Castro-Zavala A, et al. (2018) Chronic Arsenic Exposure Increases Abeta(1-42) Production and Receptor for Advanced Glycation End Products Expression in Rat Brain. Chem Res Toxicol 31(1):13–21 doi:10.1021/acs.chemrestox.7b00215

52. Nino SA, Morales-Martinez A, Chi-Ahumada E, et al. (2019) Arsenic Exposure Contributes to the Bioenergetic Damage in an Alzheimer’s Disease Model. ACS Chem Neurosci 10(1):323–336 doi:10.1021/acschemneuro.8b00278

53. Niu FW, Liu MD, Yao K, et al. (2025) Mitochondrial ROS-associated integrated stress response is involved in arsenic-induced blood-testis barrier disruption and protective effect of melatonin. Environ Int 197:109346 doi:10.1016/j.envint.2025.109346

54. O’Bryant SE, Edwards M, Menon CV, Gong G, Barber R (2011) Long-term low-level arsenic exposure is associated with poorer neuropsychological functioning: a Project FRONTIER study. Int J Environ Res Public Health 8(3):861–74 doi:10.3390/ijerph8030861

55. Pakzad D, Akbari V, Sepand MR, Aliomrani M (2021) Risk of neurodegenerative disease due to tau phosphorylation changes and arsenic exposure via drinking water. Toxicol Res (Camb) 10(2):325–333 doi:10.1093/toxres/tfab011

56. Pan H, Su Q, Hong P, et al. (2024a) Arsenic-induced mtDNA release promotes inflammatory responses through cGAS-STING signaling in chicken hepatocytes. Pestic Biochem Physiol 205:106129 doi:10.1016/j.pestbp.2024.106129

57. Pan H, Zhou L, Zou J, et al. (2024b) Arsenic trioxide induces innate immune response and inflammatory response in chicken liver via cGAS-STING/NF-kappaB pathway. Comp Biochem Physiol C Toxicol Pharmacol 286:110017 doi:10.1016/j.cbpc.2024.110017

58. Pandey KK, Mehta K, Kaur B, Dhar P (2025) Curcumin alleviates arsenic trioxide-induced neural damage in the murine striatal region. Psychopharmacology (Berl) 242(3):497–520 doi:10.1007/s00213-024-06700-y

59. Rao G, Zhong G, Hu T, et al. (2023) Arsenic Trioxide Triggers Mitochondrial Dysfunction, Oxidative Stress, and Apoptosis via Nrf 2/Caspase 3 Signaling Pathway in Heart of Ducks. Biol Trace Elem Res 201(3):1407–1417 doi:10.1007/s12011-022-03219-1

60. Rashid F, Khan KM, Saiprakash S, et al. (2025) Epidemiological Evidence on the Associations of Metal Exposure with Alzheimer’s Disease and Related Dementias Among Elderly Women. J Clin Med 14(11) doi:10.3390/jcm14113776

61. Ratnaike RN (2003) Acute and chronic arsenic toxicity. Postgrad Med J 79(933):391–6 doi:10.1136/pmj.79.933.391

62. Saggu S, Srivastava RK, McCormick L, Agarwal A, Khan MM, Athar M (2024) Neurotoxicology of warfare arsenical, diphenylarsinic acid in humans and experimental models. Chemosphere 367:143516 doi:10.1016/j.chemosphere.2024.143516

63. Shreeya T, Ansari MS, Kumar P, et al. (2023) Senescence: A DNA damage response and its role in aging and Neurodegenerative Diseases. Front Aging 4:1292053 doi:10.3389/fragi.2023.1292053

64. Sidrauski C, Acosta-Alvear D, Khoutorsky A, et al. (2013) Pharmacological brake-release of mRNA translation enhances cognitive memory. Elife 2:e00498 doi:10.7554/eLife.00498

65. Sidrauski C, McGeachy AM, Ingolia NT, Walter P (2015) The small molecule ISRIB reverses the effects of eIF2alpha phosphorylation on translation and stress granule assembly. Elife 4 doi:10.7554/eLife.05033

66. Soto C, Estrada LD (2008) Protein misfolding and neurodegeneration. Arch Neurol 65(2):184–9 doi:10.1001/archneurol.2007.56

67. Srivastava RK, Li C, Ahmad A, et al. (2016a) ATF4 regulates arsenic trioxide-mediated NADPH oxidase, ER-mitochondrial crosstalk and apoptosis. Arch Biochem Biophys 609:39–50 doi:10.1016/j.abb.2016.09.003

68. Srivastava RK, Li C, Wang Y, et al. (2016b) Activating transcription factor 4 underlies the pathogenesis of arsenic trioxide-mediated impairment of macrophage innate immune functions. Toxicol Appl Pharmacol 308:46–58 doi:10.1016/j.taap.2016.07.015

69. Sweeney P, Park H, Baumann M, et al. (2017) Protein misfolding in neurodegenerative diseases: implications and strategies. Transl Neurodegener 6:6 doi:10.1186/s40035-017-0077-5

70. Tam LM, Price NE, Wang Y (2020) Molecular Mechanisms of Arsenic-Induced Disruption of DNA Repair. Chem Res Toxicol 33(3):709–726 doi:10.1021/acs.chemrestox.9b00464

71. Thadathil N, Delotterie DF, Xiao J, Hori R, McDonald MP, Khan MM (2021) DNA Double-Strand Break Accumulation in Alzheimer’s Disease: Evidence from Experimental Models and Postmortem Human Brains. Mol Neurobiol 58(1):118–131 doi:10.1007/s12035-020-02109-8

72. Tian X, Zhang S, Zhou L, et al. (2021) Targeting the Integrated Stress Response in Cancer Therapy. Front Pharmacol 12:747837 doi:10.3389/fphar.2021.747837

73. Tian Y, Hou Q, Zhang M, Gao E, Wu Y (2025) Exposure to arsenic and cognitive impairment in children: A systematic review. PLoS One 20(2):e0319104 doi:10.1371/journal.pone.0319104

74. Tsai JC, Miller-Vedam LE, Anand AA, et al. (2018) Structure of the nucleotide exchange factor eIF2B reveals mechanism of memory-enhancing molecule. Science 359(6383) doi:10.1126/science.aaq0939

75. Tyler CR, Allan AM (2014) The Effects of Arsenic Exposure on Neurological and Cognitive Dysfunction in Human and Rodent Studies: A Review. Curr Environ Health Rep 1(2):132–147 doi:10.1007/s40572-014-0012-1

76. Vaidya N, Holla B, Heron J, et al. (2023) Neurocognitive Analysis of Low-level Arsenic Exposure and Executive Function Mediated by Brain Anomalies Among Children, Adolescents, and Young Adults in India. JAMA Netw Open 6(5):e2312810 doi:10.1001/jamanetworkopen.2023.12810

77. Wadgaonkar P, Bi Z, Wan J, et al. (2022) Arsenic Activates the ER Stress-Associated Unfolded Protein Response via the Activating Transcription Factor 6 in Human Bronchial Epithelial Cells. Biomedicines 10(5) doi:10.3390/biomedicines10050967

78. Wadgaonkar P, Chen F (2021) Connections between endoplasmic reticulum stress-associated unfolded protein response, mitochondria, and autophagy in arsenic-induced carcinogenesis. Semin Cancer Biol 76:258–266 doi:10.1016/j.semcancer.2021.04.004

79. Wang X, Huang X, Zhou L, et al. (2021) Association of arsenic exposure and cognitive impairment: A population-based cross-sectional study in China. Neurotoxicology 82:100–107 doi:10.1016/j.neuro.2020.11.009

80. Wang YC, Li X, Shen Y, et al. (2020) PERK (Protein Kinase RNA-Like ER Kinase) Branch of the Unfolded Protein Response Confers Neuroprotection in Ischemic Stroke by Suppressing Protein Synthesis. Stroke 51(5):1570–1577 doi:10.1161/STROKEAHA.120.029071

81. Watanabe R, Unuma K, Noritake K, Funakoshi T, Aki T, Uemura K (2017) Ataxia telangiectasia and rad3 related (ATR)-promyelocytic leukemia protein (PML) pathway of the DNA damage response in the brain of rats administered arsenic trioxide. J Toxicol Pathol 30(4):333–337 doi:10.1293/tox.2017-0020

82. Weng CY, Chiou SY, Wang L, Kou MC, Wang YJ, Wu MJ (2014) Arsenic trioxide induces unfolded protein response in vascular endothelial cells. Arch Toxicol 88(2):213–26 doi:10.1007/s00204-013-1101-x

83. Yan N, Li Y, Xing Y, et al. (2022) Developmental arsenic exposure impairs cognition, directly targets DNMT3A, and reduces DNA methylation. EMBO Rep 23(6):e54147 doi:10.15252/embr.202154147

84. Zhang B, Wang J, Chen J, Pan X (2025) Arsenic exposure activates microglia, inducing neuroinflammation and promoting the occurrence and development of Alzheimer’s disease-like neurodegeneration in mice. Ecotoxicol Environ Saf 297:118251 doi:10.1016/j.ecoenv.2025.118251

85. Zhang XY, Yang SM, Zhang HP, et al. (2015) Endoplasmic reticulum stress mediates the arsenic trioxide-induced apoptosis in human hepatocellular carcinoma cells. Int J Biochem Cell Biol 68:158–65 doi:10.1016/j.biocel.2015.09.009

86. Zyryanova AF, Kashiwagi K, Rato C, et al. (2021) ISRIB Blunts the Integrated Stress Response by Allosterically Antagonising the Inhibitory Effect of Phosphorylated eIF2 on eIF2B. Mol Cell 81(1):88–103 e6 doi:10.1016/j.molcel.2020.10.031

