## Supplementary file for "Arsenic Trioxide Underpins Delayed Neuroinflammation and Impaired Synaptic Integrity involving Integrative Stress Response Signaling"

**Table S1:** Primers for RT-qPCR and mitochondrial DNA copy numbers in mouse

| | | | | |
|---|---|---|---|---|
| **Il1b_mF** | tggaccttccaggatgaggaca | NM_008361.4 331 - 352 | Mouse | |
| **Il1b_mR** | gttcatctcggagcctgtagtg | NM_008361.4 478 - 457 | Mouse | 148 (with Il1b_mF) |
| **Il6_mF** | taccacttcacaagtcggaggc | NM_031168.2 216 - 237 | Mouse | |
| **Il6_mR** | ctgcaagtgcatcatcgttgttc | NM_031168.2 331 - 309 | Mouse | 116 (with Il6_mF) |
| **Il18_mF** | caaaccttccaaatcacttcct | NM_008360.2 556 - 577 | Mouse | |
| **Il18_mR** | tccttgaagttgacgcaaga | NM_008360.2 633 - 614 | Mouse | 78 (with Il18_mF) |
| **Gapdh_mR** | catcactgccacccagaagactg | NM_008084.4 606 - 628 | Mouse | |
| **Gapdh_mR** | atgccagtgagcttcccgttcag | NM_008084.4 758 - 736 | Mouse | 153 (with Gapdh_mF) |
| **mtDNA_mF1** | cgcaagggaaagatgaaagact | NC_005089.1 1214 - 1235 | Mouse | |
| **mtDNA_mR1** | tcgtttggtttcggggtttc | NC_005089.1 1345 - 1326 | Mouse | 132 (with mtDNA_mF1) |
| **nDNA_mF1** | acagacagacagacagacattt | NC_000072 82737656 - 7682 | Mouse | |
| **nDNA_mR1** | agaggagggaggccatgag | NC_000072 82737795 - 7777 | Mouse | 140 (with nDNA_mF1) |

**Table S2:** Antibodies for Western blot

| S/NO | Antibody | Manufacturer & location | catalog number | Dilution |
|---|---|---|---|---|
| 1 | p-PERK | Cell Signaling, USA | 3179 | 1:1000 |
| 2 | p-eIF2α (Ser51) | Cell Signaling, USA | 3398 | 1:1000 |
| 3 | CHOP | ProteinTech, USA | 15204-1-AP | 1:1000 |
| 4 | ATF4 | Invitrogen | MA5-32364 | 1:1000 |
| 5 | γ-H2A.X (Ser139) | Abcam, USA | ab11174 | 1:500 |
| 6 | STING | Cell Signaling, USA | 13647 | 1:500 |
| 7 | STING | ProteinTech, USA | 19851-1-AP | 1:250 |
| 8 | Caspase-1 | Santa Cruz Biotechnology | sc-392736 | 1:1000 |
| 9 | NLRP3 | Adipogen Corporation, USA | AG-20B-0014-C100 | 1:1000 |
| 10 | β-actin | Invitrogen | MA1-140 | 1:1000 |
| 11 | Iba1 | Synaptic System USA | 234 009 | 1:500 |
| 12 | MAP2 | EnCor Biotechnology, USA | CPCA-MAP2 | 1:1000 |
| 13 | PSD95 (D27E11) | Cell Signaling, USA | 3450 | 1:1000 |
| 14 | p16 INK4A (F2T7H) | Cell Signaling, USA | 23200 | 1:1000 |

**Fig. S1**

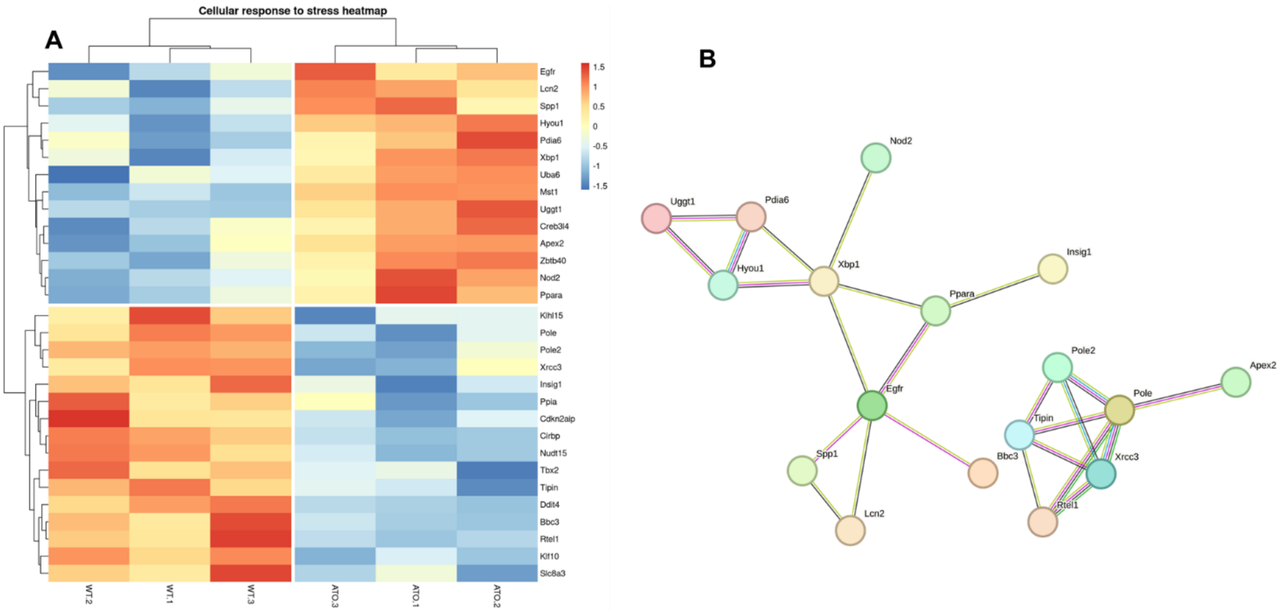
A
Cellular response to stress heatmap
Egfr
Lcn2
Spp1
Hyou1
Pdia6
Xbp1
Uba6
Mst1
Uggt1
Creb3l4
Apex2
Zbtb40
Nod2
Ppara
Klhl15
Pole
Pole2
Xrcc3
Insig1
Ppia
Cdkn2aip
Cirbp
Nudt15
Tbx2
Tipin
Ddit4
Bbc3
Rtel1
Klf10
Slc8a3
1.5
1
0.5
0
-0.5
-1
-1.5
WT.2
WT.1
WT.3
ATO.3
ATO.1
ATO.2
B
Nod2
Uggt1
Pdia6
Hyou1
Xbp1
Insig1
Ppara
Egfr
Spp1
Lcn2
Bbc3
Pole2
Apex2
Pole
Tipin
Xrcc3
Rtel1

**Fig. S2**

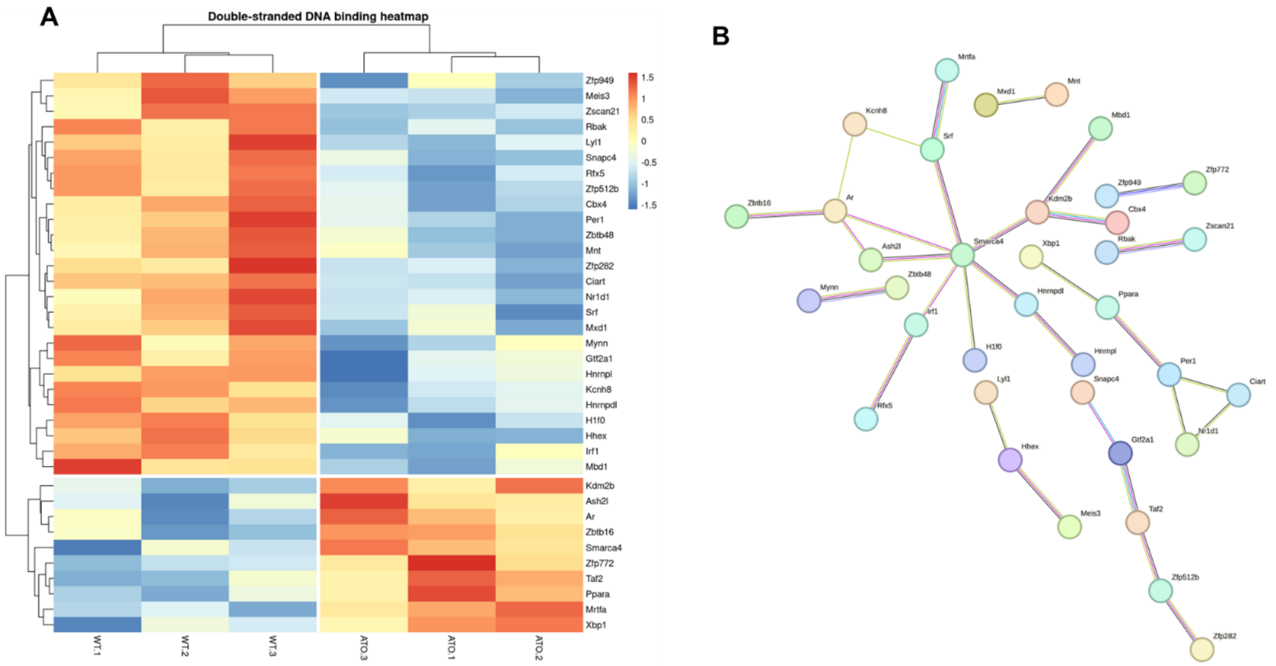
A
Double-stranded DNA binding heatmap
Zfp949
Meis3
Zscan21
Rbak
Lyl1
Snapc4
Rfx5
Zfp512b
Cbx4
Per1
Zbtb48
Mnt
Zfp282
Ciart
Nr1d1
Srf
Mxd1
Mynn
Gtf2a1
Hnrnpl
Kcnh8
Hnrnpdl
H1f0
Hhex
Irf1
Mbd1
Kdm2b
Ash2l
Ar
Zbtb16
Smarca4
Zfp772
Taf2
Ppara
Mrtfa
Xbp1
1.5
1
0.5
0
-0.5
-1
-1.5
WT.1
WT.2
WT.3
ATO.3
ATO.1
ATO.2
B
Mrtfa
Mxd1
Mnt
Kcnh8
Srf
Mbd1
Zfp949
Zfp772
Zbtb16
Ar
Kdm2b
Cbx4
Zscan21
Smarca4
Rbak
Xbp1
Ash2l
Zbtb48
Mynn
Hnrnpdl
Ppara
Irf1
H1f0
Hnrnpl
Per1
Ciart
Lyl1
Snapc4
Rfx5
Nr1d1
Gtf2a1
Hhex
Meis3
Taf2
Zfp512b
Zfp282

**Fig. S3**

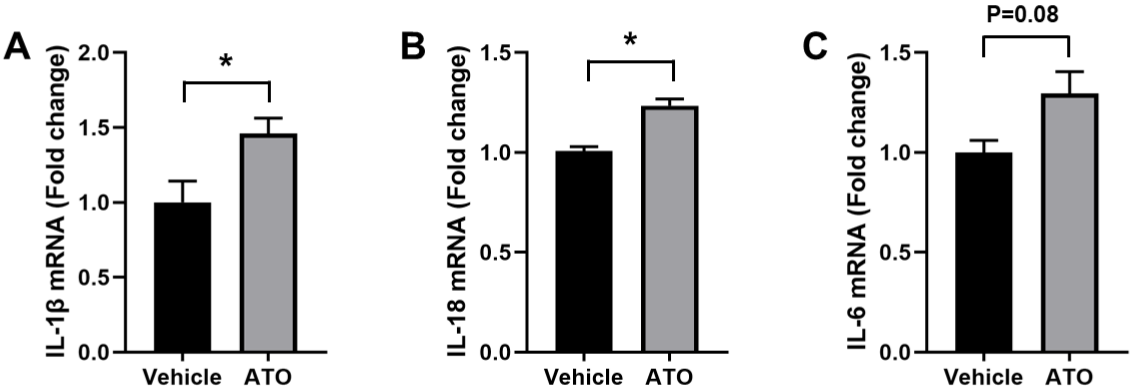
A
IL-1β mRNA (Fold change)
2.0
1.5
1.0
0.5
0.0
*
Vehicle
ATO
B
IL-18 mRNA (Fold change)
1.5
1.0
0.5
0.0
*
Vehicle
ATO
C
IL-6 mRNA (Fold change)
P=0.08
1.5
1.0
0.5
0.0
Vehicle
ATO

**Fig. S4**

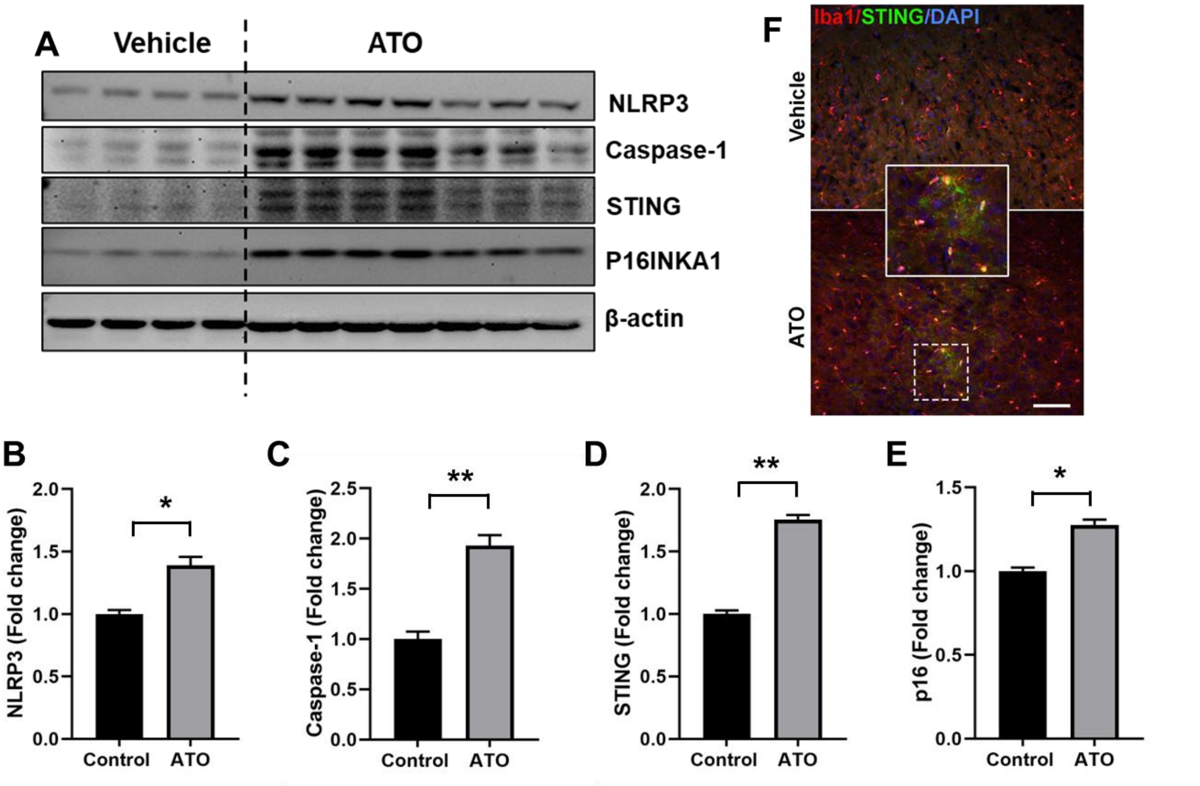
A
Vehicle
ATO
NLRP3
Caspase-1
STING
P16INKA1
β-actin
F
Iba1/STING/DAPI
Vehicle
ATO
B
NLRP3 (Fold change)
2.0
1.5
1.0
0.5
0.0
*
Control
ATO
C
Caspase-1 (Fold change)
2.5
2.0
1.5
1.0
0.5
0.0
**
Control
ATO
D
STING (Fold change)
2.0
1.5
1.0
0.5
0.0
**
Control
ATO
E
p16 (Fold change)
1.5
1.0
0.5
0.0
*
Control
ATO

**Figure legends**

**Fig. S1**. Stress response and protein-protein interaction (PPI) networks in WT mice following vehicle or ATO exposure. (A) Heat map showing differential expression of stress response-related genes between vehicle-treated and ATO-exposed WT mice. (B) PPI network highlighting major hub proteins involved in the stress response pathway.

**Fig. S2.** DNA double-strand binding and PPI networks in WT mice following vehicle or ATO exposure. (A) Heat map showing differential expression of DNA double-strand repair-related genes between vehicle-treated and ATO-exposed WT mice. (B) PPI network highlighting key hub proteins in the DNA double-strand repair pathway.

**Fig. S3.** Induction of pro-inflammatory cytokines in WT mice following ATO exposure. Relative mRNA expression levels of IL-1β, IL-18, and IL-6 were measured in the hippocampus of WT mice 4 weeks after ATO treatment compared with vehicle controls by RT-qPCR. N=4 mice/group Data were analyzed using an unpaired t-test and are presented as mean ± SEM. $^{*}P < 0.05$.

**Fig. S4.** ATO exposure induces neuroinflammation in the cortex of WT mice. (A-D). Representative immunoblots and quantification of STING, NLRP3, caspase-1, and p16 protein levels in the cortex of WT mice treated with vehicle or ATO. N=4-7 mice/. Data are presented as mean ± SEM and compared between groups using a two-tailed t-test. $^{*}P < 0.05$; $^{**}P < 0.01$. (E, F) Representative images of cortical brain sections showing STING expression in microglia in WT mice exposed to vehicle or ATO.
